## supplemental figures for "Styxl2 regulates *de novo* sarcomere assembly by binding to non-muscle myosin IIs and promoting their degradation"

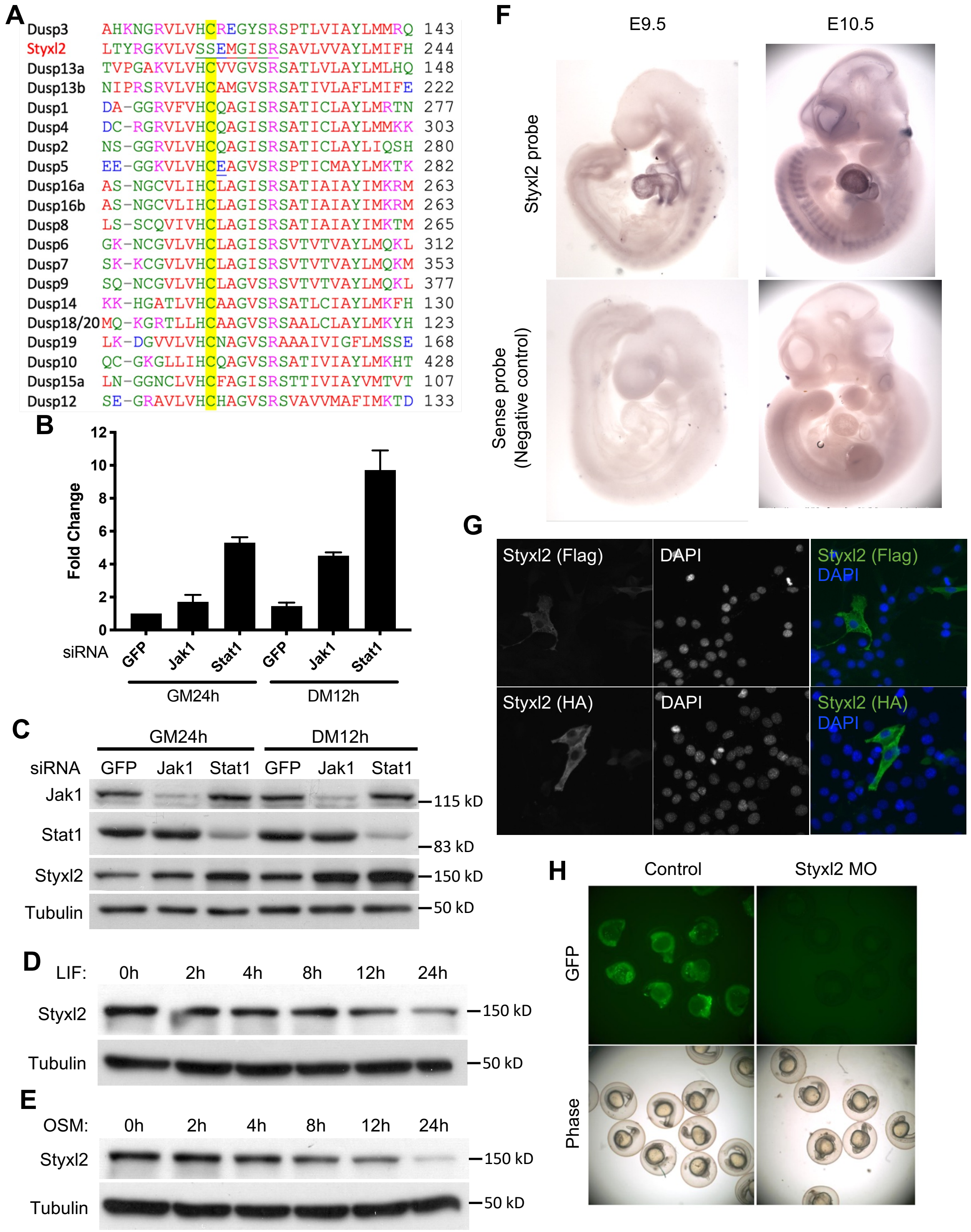


**Figure 1-figure supplement 1. Styxl2 was downstream target of Jak1-Stat1 pathway.** (A) Multiple sequence alignment of the region surrounding the active motif of the DSPc domain among mouse dual-specificity phosphatases and Styxl2. The conserved Cys in functional phosphatases was highlighted in yellow. (B, C) C2C12 cells were transfected with various siRNAs as indicated. After growing in GM for 24 hours (h), cells were harvested either right away (GM 24h) or after growing in DM for an additional 12 h (DM 12h). The total RNA and soluble whole cell lysates were subjected to RT-qPCR (B) or Western blot analysis (C), respectively. (D, E) C2C12 cells were treated with 10 ng/ml of LIF or OSM for various times. Cell extracts were subjected to Western blot analysis. (F) Mouse embryos at E9.5 and E10.5 were subjected to *in situ* hybridization using either sense (control) or antisense probe specific for mouse *Styxl2*. (G) C2C12 cells were transfected with constructs expressing Flag- or HA-tagged Styxl2 for 24 h. Cells were then fixed and subjected to immunostaining for Flag or HA. (H) Zebrafish zygotes were co-injected with a plasmid encoding GFP fused with the sequence targeted by *Styxl2*-MO at the start codon together with or without *Styxl2*-MO. The phase-contrast and fluorescent images were taken and the representative images were shown.

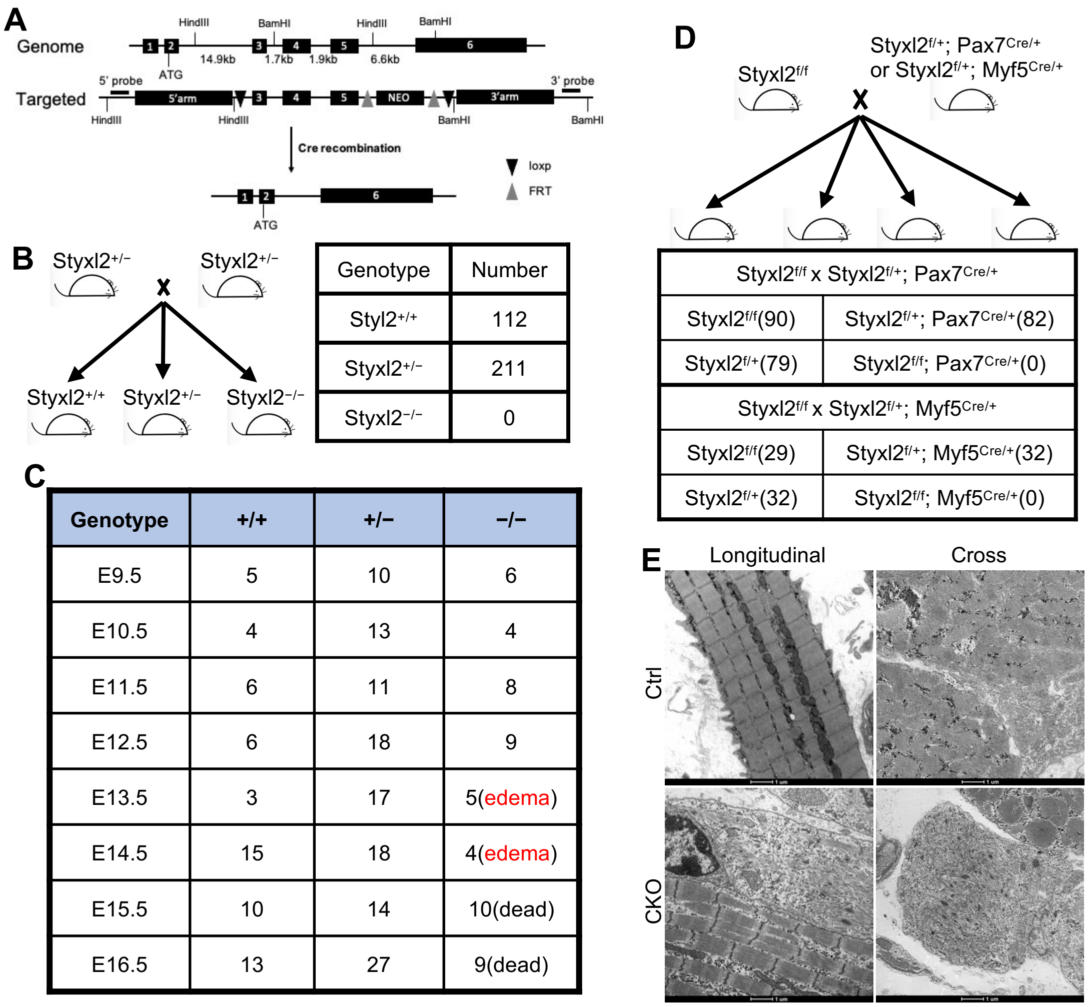


**Figure 2-figure supplement 1. Both germline and conditional deletion of *Styxl2* caused lethality.** (A) The schematic of the genomic locus of mouse *Styxl2* and the targeting strategy. The probes used for Southern blot are indicated by black bars. (B, C) The schematic of the mating strategy to generate *Styxl2* germ-line KO mice was shown (B). The number of live pups (B) and that of embryos (C) of different genotypes at different embryonic stages were counted and presented. (D) The schematic of the mating strategy to generate conditional *Styxl2* KO mice and the number of live pups (in brackets) older than 1 day were shown. (E) Hindlimb muscle sections from control (Ctrl) and *Styxl2* CKO P1 mice were subjected to electron microscopy analysis. Scale bar, 1 μm.

**A**

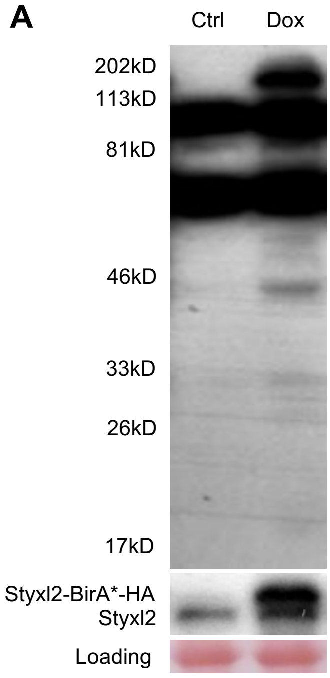


**B**

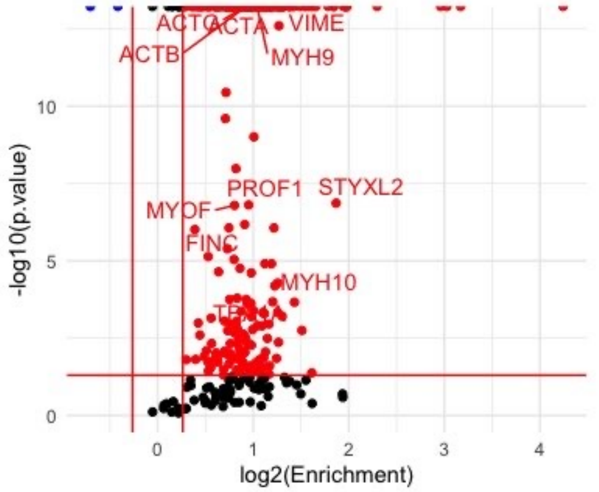


**C**

| Accession | Enrichment | Peptide No. |
|---|---|---|
| MYH9 | 2.057 | 75 |
| ACTB | 1.902 | 71 |
| ACTG | 1.895 | 66 |
| ACTA | 1.967 | 36 |
| VIME | 2.41 | 27 |
| STYXL2 | 3.65 | 24 |
| PROF1 | 1.935 | 16 |
| TBA1A | 1.662 | 13 |
| MYH10 | 2.285 | 12 |
| MYOF | 1.746 | 12 |
| FINC | 1.741 | 11 |

**Figure 4-figure supplement 1. Identification of Styxl2-interacting partners by BioID.** (A) A representative gel of samples from BioID before mass spectrometry analysis. Whole cell lysates from control (Ctrl) and Doxycycline-treated (Dox) cells were subjected to Western blot using Streptavidin-HRP (top panel) or an anti-Styxl2 antibody (middle panel). (B) The biotinylated proteins identified by mass spectrometry were plotted with the $\log_2$Enrichment scores on the X-axis and the minus $\log_{10}$(p.value) on the Y-axis. The Enrichment score denotes the ratio of the peak intensity of reporter ions. Red dots indicate proteins with the Enrichment score > 1.2 and p value < 0.05. Blue dots indicate proteins with the Enrichment score < 0.83 and p value < 0.05. (C) Selected candidate proteins with more than 10 recovered peptides were shown.

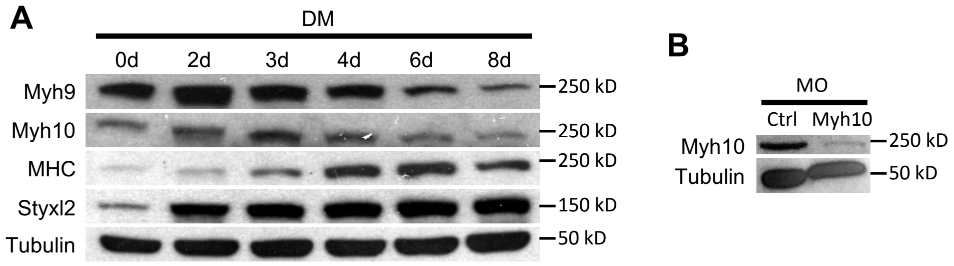


**Figure 5-figure supplement 1.** (A) Confluent C2C12 cells were induced to differentiate in DM and harvested at different time points. Soluble whole cell lysates were subjected to Western blot analysis. (B) Zebrafish zygotes were injected with control (Ctrl) or Myh10-MO and the fish were collected for Western blot analysis at 48 hpf.

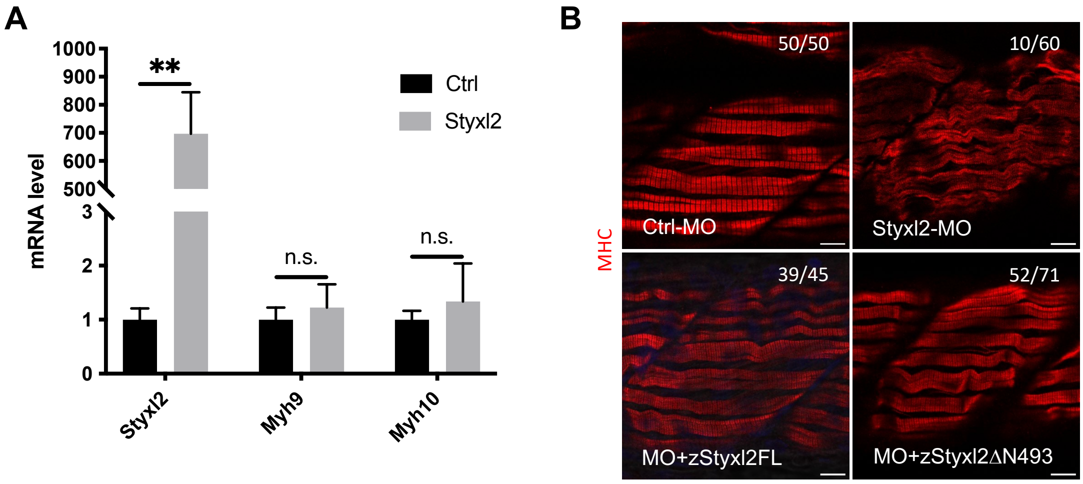


**Figure 6-figure supplement 1.** (A) Control (Ctrl) or Styxl2 DNA constructs were transfected into C2C12 cells. 24 h later, the mRNA levels of different genes were determined by qPCR. **, p-value <0.01. n.s.: not significant. (B) Zebrafish zygotes were injected with various morpholinos as indicated with or without the co-injected *Styxl2* mRNAs. z*Styxl2*FL: the full-length fish *Styxl2* mRNA; z*Styxl2*ΔN493: the truncated fish *Styxl2* mRNA missing the 5' region encoding the N-terminal 493 aa. Representative immunofluorescent images of myosin heavy chain (MHC) were shown. The number in numerator at the top right corner of each image represents fish embryos showing normal MHC staining pattern, while that in denominator represents the total number of embryos examined. Scale bar: 10 [illegible]m.

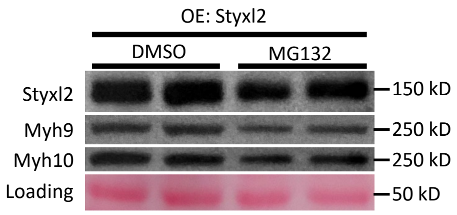


**Figure 7-figure supplement 1.** C2C12 cells were transfected with a Styxl2-expressing plasmid. 12 hours later, MG132 (4 □M) or DMSO was added into the medium. After another 12 hours, the cells were harvested for Western blot analysis. OE, overexpression.
